# Gut microbiota composition related with *Clostridium difficile*-positive diarrhea and *C. difficile* type (A^+^B^+^, A^-^B^+^, and A^-^B^-^) in ICU hospitalized patients

**DOI:** 10.1101/126995

**Authors:** Anhua Wu, Juping Duan, Sidi Liu, Xiujuan Meng, Pengcheng Zhou, Qingya Dou, Chunhui Li

## Abstract

**Background:** Gut microbiota composition of intensive care unit (ICU) patients suffering from *Clostridium difficile*-positive diarrhea (CDpD) is still poorly understood. This study aims to use 16S rDNA (and metagenome) sequencing to compare the microbiota composition of 58 (and 5) ICU patients with CDpD (CDpD group), 33 (and 4) ICU patients with *C. difficile* negative diarrhea (CDnD group), and 21 (and 5) healthy control subjects (control group), as well as CDpD patients in A^+^B^+^ (N=34; A/B: *C. difficile TcdA/B*), A^-^B^+^ (N=7), and A^-^B^-^ (N=17) subgroups. For 16S rDNA data, OTU clustering (tool: UPARSE), taxonomic assignment (tool: RDP classifier), α-diversity and β-diversity analyses (tool: QIIME) were conducted. For metagenome data, metagenome assembly (tool: SOAP), gene calling (tools: MetaGeneMark, CD-HIT, and SoapAligner), unigene alignment (tool: DIAMOND), taxon difference analysis (tool: Metastats), and gene annotation (tool: DIAMOND) were performed.

**Results:** The microbial diversity of CDpD group was lower than that of CDnD and control groups. The abundances of 10 taxa (e.g. Deferribacteres, Cryptomycota, Acetothermia) in CDpD group were significantly higher than that in CDnD group. The abundances of Saccharomycetes and Clostridia were significantly lower in CDpD in comparison with control. A^+^B^+^, A^-^B^+^ and A^-^B^-^ subgroups couldn’t be separated in principal component analysis, while some taxa are significantly different between A^+^B^+^ and A^-^B^-^ subgroups.

**Conclusion:** CDpD might relate to the decrease of beneficial taxa (i.e. Saccharomycetes and Clostridia) and the increase of harmful taxa (e.g. Deferribacteres, Cryptomycota, Acetothermia) in gut microbiota in ICU patients. *C. difficile* type might be slightly associated with gut microbiota composition.

## Introduction

As a result of infection prior admission, illness-associated immunosuppression, and invasive devices in intensive care unit (ICU), microbial infection is frequently developed in patients in general ICU (Vincent et al. 2016). Thus, antibiotics are often used in ICU patients (Vincent et al. 2016), including Meropenem, Piperacillin/tazobactam, Imipenem/Cilastatin, Cefuroxime, Ceftriaxone, Ceftazidime, Cefepime, Cefperazone-Sulbactam, Latamoxef, etc. However, antibiotic treatment is the most crucial risk factor for the infection with *Clostridium difficile* (also known as *Peptoclostridium difficile*) (Stevens et al. 2011; Yutin and Galperin 2013), as it can adversely affect gut indigenous microbiota composition and decrease the colonization resistance to *C. difficile* (Britton and Young 2012). Consequently, *C. difficile*-associated diarrhea happens in 9.45 cases per 10,000 patient-days in ICU (Lee et al. 2015).

Nevertheless, current antibiotic therapies for *C. difficile* infection (CDI) like vancomycin have limited efficacy (Bagdasarian et al. 2015), and fecal microbiota transplantation (FMT) is recommended as an alternative therapy in curing highly-recurrent CDI failed to respond to vancomycin treatment (Kelly et al. 2015). The success of FMT emphasizes the importance of understanding gut microbiome in CDI patients. Therefore, a better understanding of the relationships between microbiota, *C. difficile*, and diarrhea in ICU patients may contribute to identifying novel therapeutic targets for diarrhea in ICU patients.

In recent years, high-throughput deep-sequencing of 16S rDNA and metagenome has been applied to investigating microbiota composition. Milani et al. used 16S rDNA and metagenome sequencing to study the gut microbiota compositions of three groups of elderly (age□≥□65) hospitalized patients, involving 30 CDI-negative patients not exposed to antibiotics (AB^-^ group), 29 CDI-negative patients exposed to antibiotics (AB^+^ group), and 25 CDI-positive patients (CDI group) (Milani et al. 2016). The microbial diversity of CDI-positive group was significantly lower than that of CDI-negative group. CDI was found to relate with decrease in gut commensals like *Bacteroides*, *Alistipes*, *Lachnospira*, and *Barnesiella*, while antibiotic treatment in CDI-negative patients might associate with the depletion of commensals like *Alistipes* (Milani et al. 2016). Schubert et al. utilized 16S rDNA sequencing to study the gut microbiota compositions of CDI cases, diarrheal controls, and non-diarrheal controls (Schubert et al. 2014). Statistical models were developed for CDI and diarrhea by incorporating clinical and demographic data with microbiome data, and loss of several species in Ruminococcaceae, Lachnospiraceae, *Bacteroides*, and Porphyromonadaceae might associate with CDI (Schubert et al. 2014). Generally, only *C. difficile* that produces toxin A (an intestinotoxin, *TcdA*) and/or toxin B (a cytotoxin, *TcdB*) can cause gastrointestinal diseases in humans. However, precise microbiome alterations related with diarrhea and *C.* difficile-positive diarrhea (CDpD) have not been completely elucidated in ICU patients. Also, relationships between *C. difficile* type (i.e. A^+^B^+^, A^-^B^+^, and A^-^B^-^) and gut microbiota composition have not been investigated.

In this study, we utilized 16S rRNA and metagenome deep-sequencing to characterize the gut microbiota composition in healthy subjects and ICU patients stratified for *C. difficile*, as well as the gut microbiota composition in ICU patients with A^+^B^+^ CDpD, A^-^B^+^ CDpD, and A^-^B^-^ CDpD. Our results may help to understand the mechanisms underlying diarrhea and CDpD development in ICU patients and identify potential candidates for curative or preventive microbe therapy.

## Results

### Clinical features of subjects

For Result-1, subjects in CDpD group (N=58), CDnD group (N=33), and control group (N=21) were similar for gender, and subjects in CDpD and CDnD groups were similar for age, gender, inpatient days, diabetes, cancer, hematopathy, respiratory failure, renal insufficiency, tuberculosis, surgery, and exposure to immunosuppressant and glucocorticoids (Table 1).

**Table 1.**
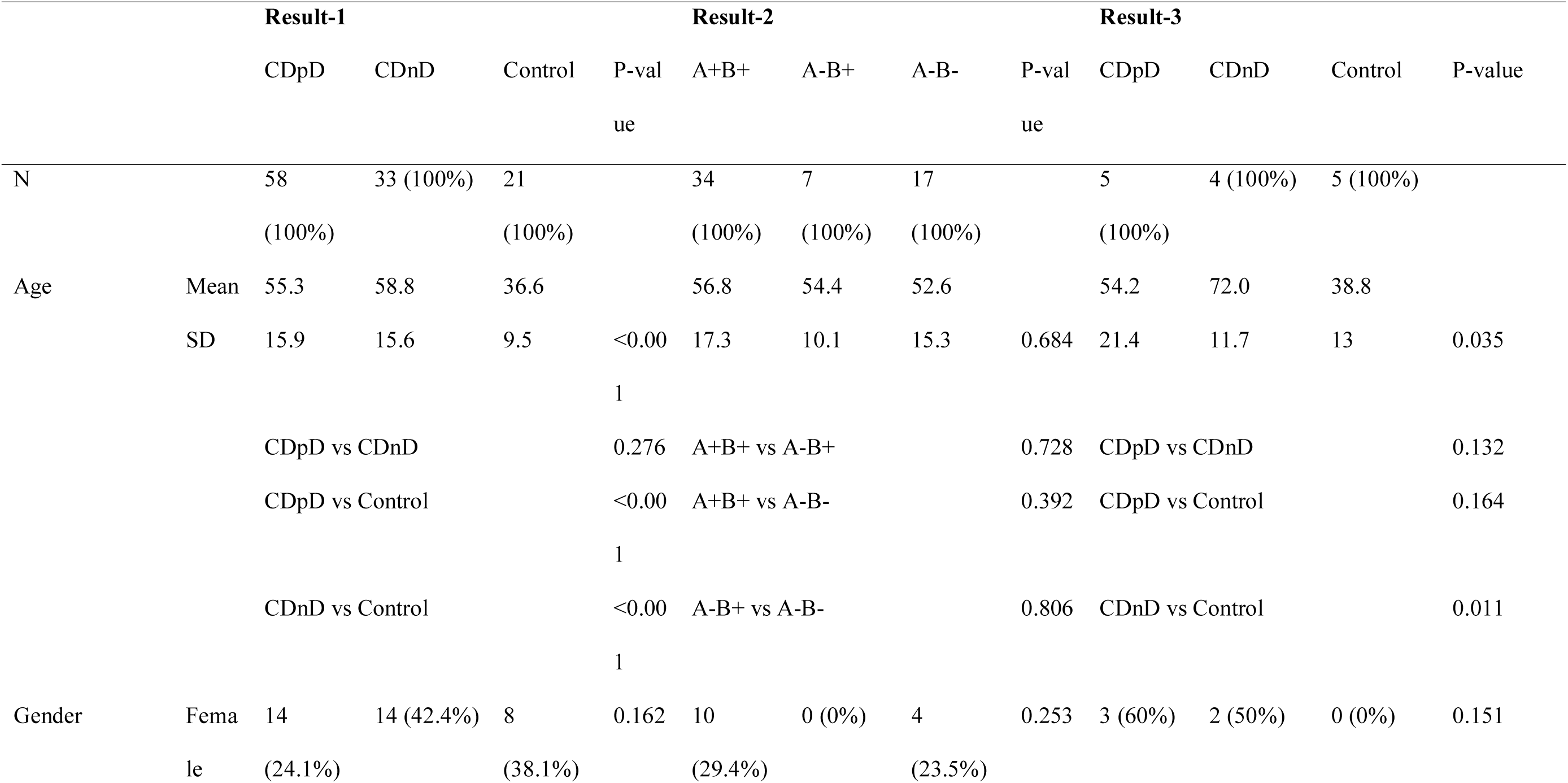
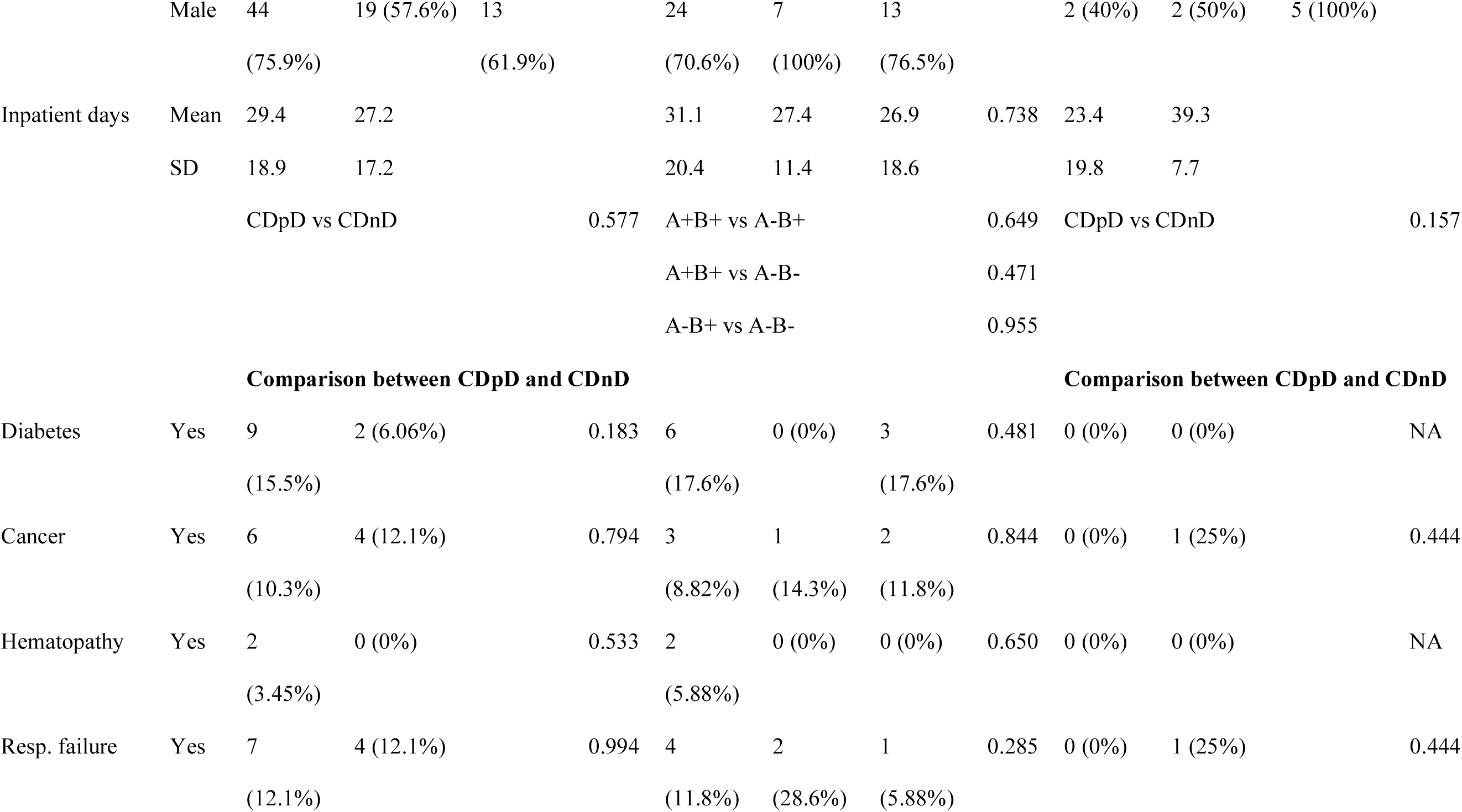
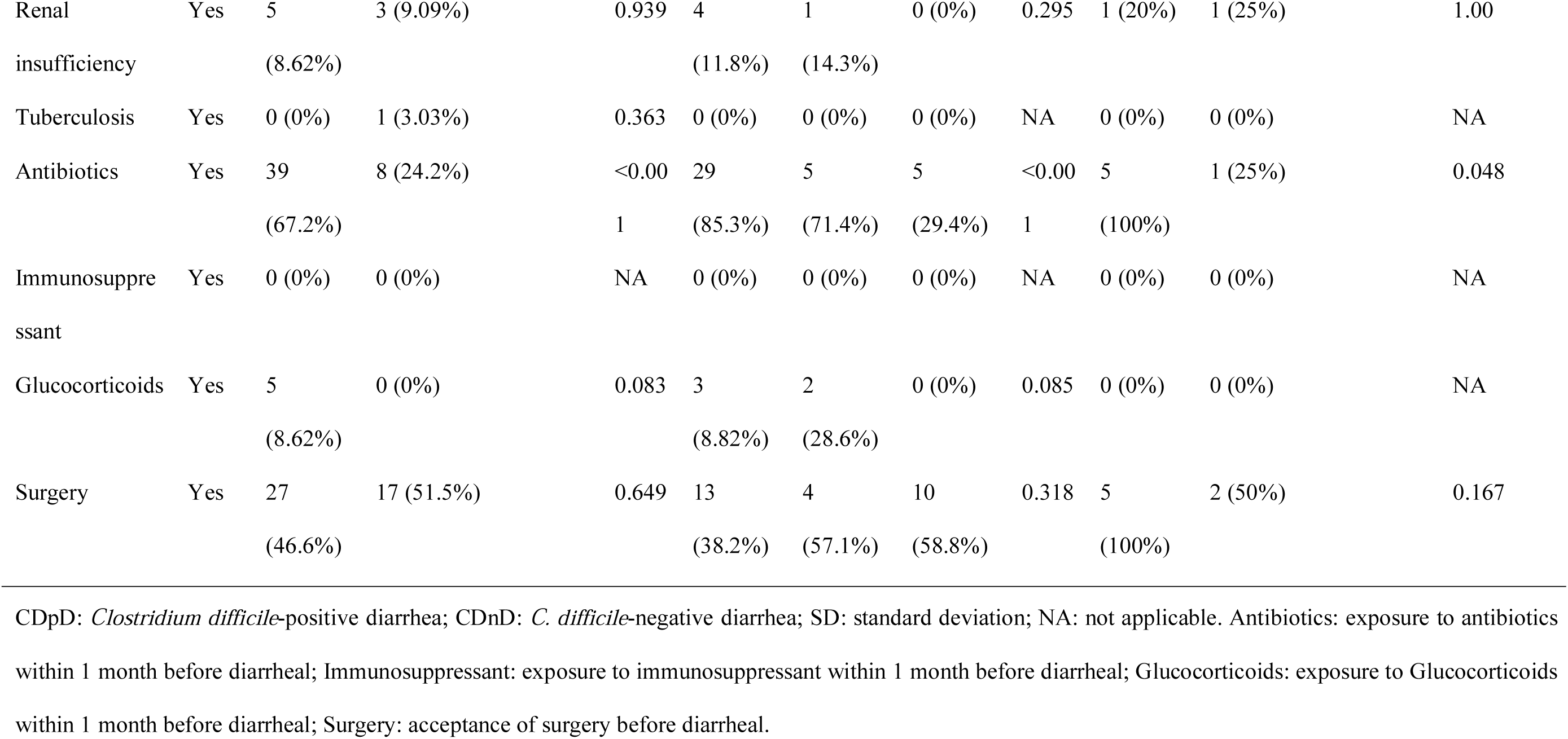
Demographic and clinical characteristics of the population in each experimental group

For Result-2, subjects in A^+^B^+^ subgroup (N=34), A^-^B^+^ subgroup (N=7), and A^-^B^-^ subgroup (N=17) of CDpD group were similar for age, gender, inpatient days, diabetes, cancer, hematopathy, respiratory failure, renal insufficiency, tuberculosis, surgery, and exposure to immunosuppressant and glucocorticoids (Table 1).

For Result-3, subjects in CDpD group (N=5), CDnD group (N=4), and control group (N=5) were similar for gender, and subjects in CDpD and CDnD groups were similar for age, gender, inpatient days, diabetes, cancer, hematopathy, respiratory failure, renal insufficiency, tuberculosis, surgery, and exposure to immunosuppressant and glucocorticoids (Table 1).

### Analysis of CDpD, CDnD, and control groups

#### α-diversity (16S rDNA sequencing data)

Chao1 index, Shannon index, and number of observed species were assessed.. In Result-1, Chao1 index (i.e. community richness) in CDpD group was lower than that in CDnD and control groups (Figure 1A); Shannon index (i.e. community diversity) in CDpD group was similar to that in CDnD group but much lower than that in control group (Figure 1B); observed species in CDpD group were much less than that in CDnD and control groups (Figure 1C).

**Figure 1.**
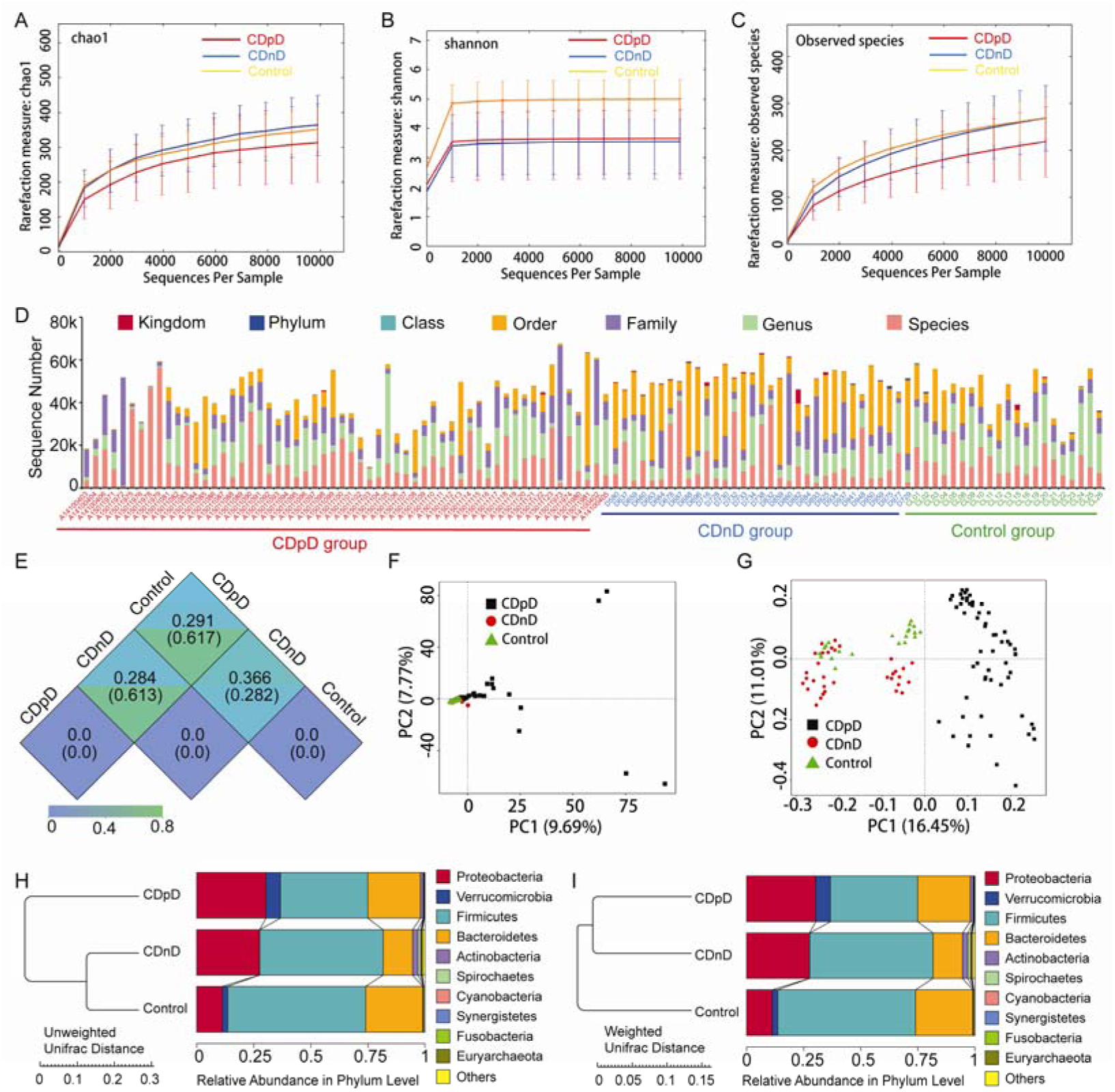
α-diversity and β-diversity of CDpD, CDnD, and control samples based on 16S rDNA sequencing data. (A) Rarefaction-curve based on Chao1 index. (B) Rarefaction-curve based on Shannon index. (C) Rarefaction-curve based on observed species. (D) Distribution of sequences in Kingdom, Phylum, Class, Order, Family, Genus, and Species. (E) Unifrac distance between groups. Upper number: weighted unifrac distance; Lower number: un-weighted unifrac distance. Both numbers represent the index for differences of taxon-diversity between groups. (F) PCA based on OTUs. (G) PCoA based on un-weighted unifrac distance. (H) Sample clustering based on un-weighted unifrac distance and relative abundance at Phylum level. (I) Sample clustering based on weighted unifrac distance and relative abundance at Phylum level. CDpD: *Clostridium difficile*-positive diarrhea; CDnD: *C. difficile*-negative diarrhea; PCA: principal component analysis; OTUs: Operational Taxonomic Units; PCoA: principal co-ordinates analysis.

#### Core-pan gene analysis (metagenome sequencing data)

In Result-3, number of non-redundant genes declined along with the increase in sample number in core-gene curve (Figure 2A), whereas number of non-redundant genes increased along with the increase in sample number in pan-gene curve (Figure 2B). These results supported a further detailed analysis on metagenome sequencing data.

**Figure 2.**
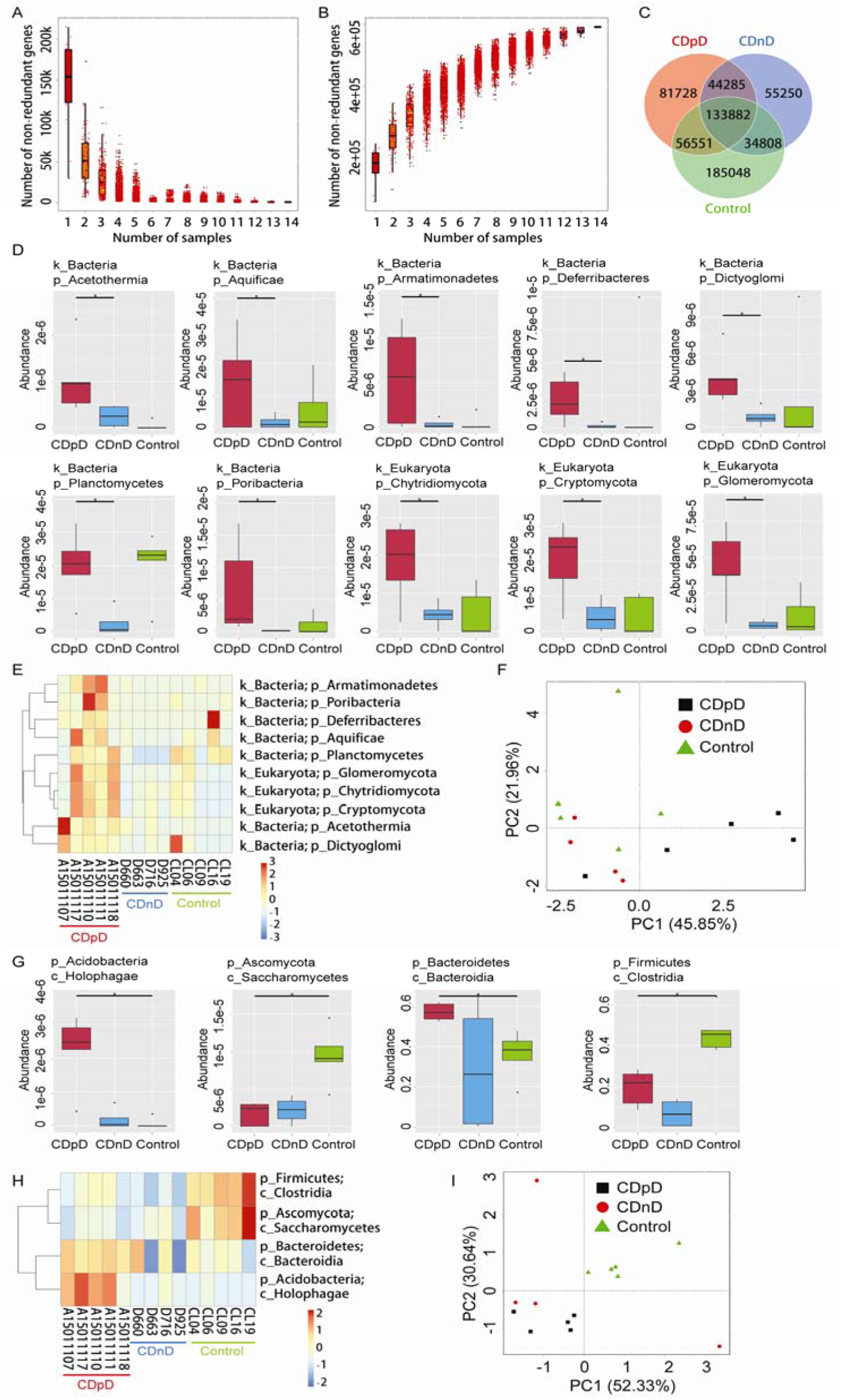
Gene prediction and taxonomic annotation based on metagenome sequencing data. (A) Rarefaction-curve based on core-gene. (B) Rarefaction-curve based on pan-gene. (C) Venn analysis of non-redundant genes. (D) Significant taxonomic differences between groups (Phylum-level). (E) Clustering based on the significant taxonomic differences at Phylum level. (F) PCA based on the significant taxonomic differences at Phylum level. (G) Significant taxonomic differences between groups (Class-level). (H) Clustering based on the significant taxonomic differences at Class level. (I) PCA based on the significant taxonomic differences at Class level. PCA: principal component analysis; CDpD: *Clostridium difficile*-positive diarrhea; CDnD: *C. difficile*-negative diarrhea.

#### OTUs and predicted genes in each group

In Result-1, 2125 16S rDNA-based OTUs were clustered, among which 513 OTUs were shared by the three groups. In Result-3, 591552 genes were predicted based on genome sequence (Figure 2C).

#### Taxonomic annotation (16S rDNA sequencing data)

The distribution of sequences in Kingdom, Phylum, Class, Order, Family, Genus, and Species were investigated. Sequences assigned into Order in CDnD group were much more than that in CDpD and control groups (Figure 1D).

#### β-diversity (16S rDNA sequencing data)

In Result-1, the un-weighted unifrac distances were 0.282, 0.617, and 0.61 between CDnD and control, between CDpD and control, and between CDpD and CDnD (Figure 1D). PCA was performed based on OTUs, and the majority of CDpD samples could be distinguished from controls despite the high inter-sample variability (Figure 1E). PCoA was conducted based on un-weighted unifrac distance, and CDpD samples could be clearly separated from CDnD samples and controls (Figure 1F). Besides, CDpD, CDnD, and control groups could be distinguished from each other based on un-weighted/weighted unifrac distance and relative abundance at Phylum level (Figure 1G and 1H).

#### Differences of community composition between groups

Relative abundances of top 10 taxa in Phylum, Class, Order, Family, and Genus were studied. In Result-1, compared with CDnD and control groups, relative abundance of Firmicutes was lower, whereas relative abundances of Proteobacteria, Verrucomicrobia, Verrucomicrobiae, Gammaproteobacteria, Verrucomicrobiales, Enterobacteriales, Porphyromonadaceae, Verrucomicrobiaceae, Enterobacteriaceae, *Parabacteroides*, and *Akkermansia* were higher in CDpD group (Figure 1H and 3A-D).

**Figure 3.**
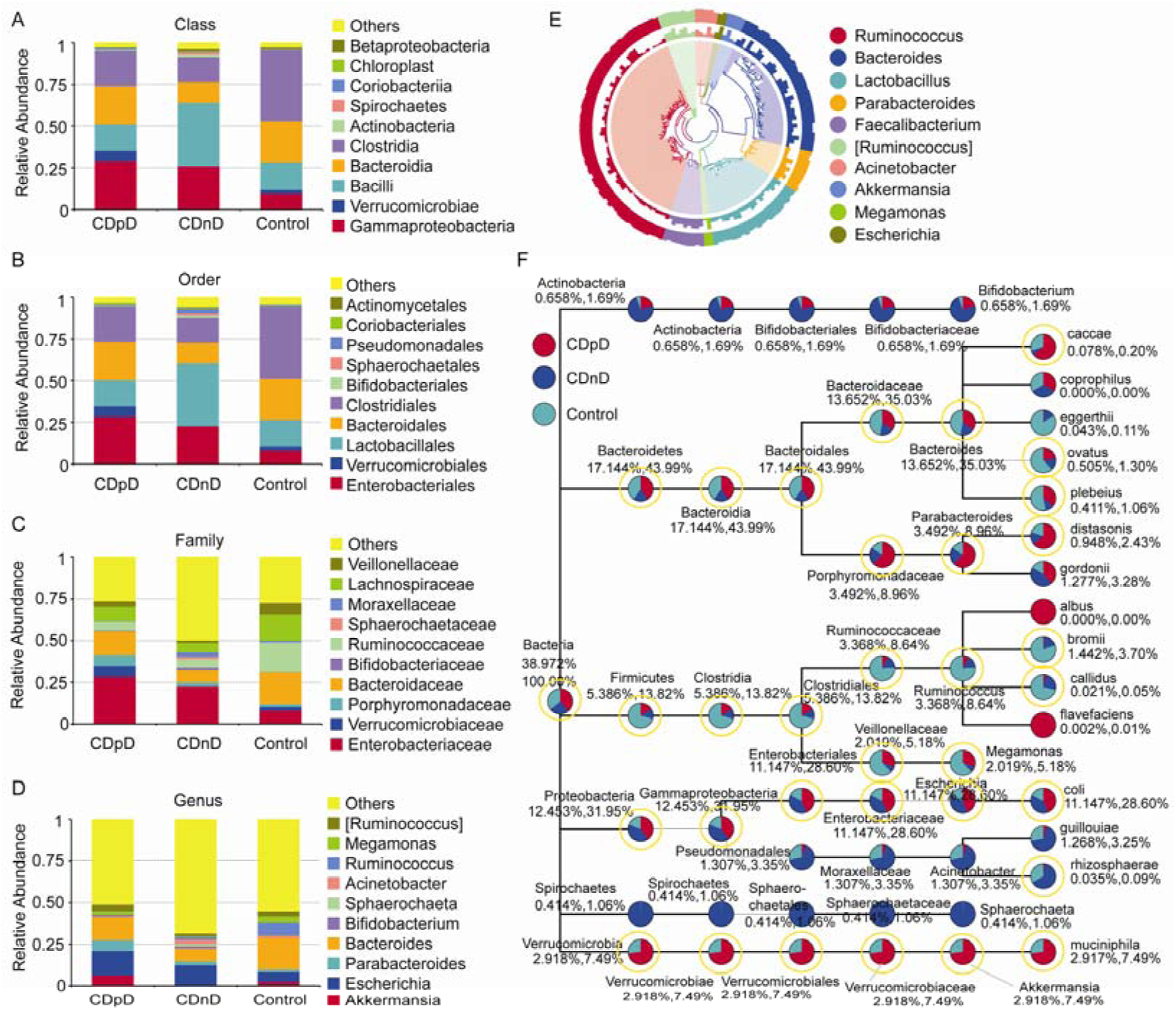
Taxonomic differences and phylogenetic relationships based on 16S rDNA sequencing data in CDpD, CDnD, and control groups. (A) Taxonomic differences between CDpD, CDnD, and control groups (Class-level). (B) Taxonomic differences between CDpD, CDnD, and control groups (Order-level). (C) Taxonomic differences between CDpD, CDnD, and control groups (Family-level). (D) Taxonomic differences between CDpD, CDnD, and control groups (Genus-level). (E) Phylogenetic relationships between OTUs annotated into top 10 Genera. Inner cycle: phylogenetic tree based on representative sequences of OTUs; Middle cycle: relative abundances of OTUs; Outer cycle: reliability of OUT annotation. (F) Taxonomic differences in selected taxa at the levels of Phylum, Class, Order, Family, and Genus. The former number = (mean relative abundance of this taxon in all sample)/(mean relative abundance of all taxa in all sample)×100%; The later number = (mean relative abundance of this taxon in all sample)/(mean relative abundance of selected taxa in all sample)×100%. Taxon with yellow cycle: this taxon was balanced across sample. CDpD: *Clostridium difficile*-positive diarrhea; CDnD: *C. difficile*-negative diarrhea; OTUs: Operational Taxonomic Units.

In Result-3, significant differences were identified between CDpD and CDnD groups in the abundances of 10 taxa (Dictyoglomi, Acetothermia, Cryptomycota, Chytridiomycota, Glomeromycota, Planctomycetes, Aquificae, Deferribacteres, Poribacteria, and Armatimonadetes; Figure 2D-E), based on which the majority of CDpD samples could be distinguished from CDnD samples in PCA in spite of high inter-sample variability (Figure 2F). Also, significant differences were found between CDpD and control groups in the abundances of Holophagae, Bacteroidia, Saccharomycetes, and Clostridia (Figure 2G-H), based on which CDpD samples could be clearly distinguished from controls in PCA (Figure 2I). In both Result-1 and Result-3, the relative abundance of Clostridia in CDpD group was lower than that in control group.

#### Phylogenetic relationship (16S rDNA sequencing data)

Phylogenetic relationships between the representative sequences of all OTUs corresponding to top 10 Genera are shown in Figure 3E. Besides, a taxonomic tree was constructed, showing the taxonomic differences in selected taxa at the levels of Phylum, Class, Order, Family, and Genus. To wipe out the influences of sample bias, we marked the taxa which were balanced across sample. In these taxa, the abundances of Porphyromonadaceae, *Parabacteroides*, *Parabacteroides distasonis*, and *Bacteroides caccae* in CDpD group were much higher than that in CDnD and control groups. These taxa might be associated with CDpD. Firmicutes, Veillonellaceae, *Megamonas*, Clostridia, Clostridiales, Ruminococcaceae, *Ruminococcus*, *Ruminococcus Bromii*, and *Ruminococcus callidus* in control group were much higher than that in CDnD and CDpD groups. These taxa might be associated with diarrhea. The abundances of Verrucomicrobia, Verrucomicrobiae, Verrucomicrobiales, Verrucomicrobiaceae, and *Akkermansia muciniphila* in CDnD group were much lower than that in CDpD and control groups. These taxa might be associated with CDnD.

#### Pathway annotation (metagenome sequencing data)

Based on KEGG database, most of the predicted genes related with Metabolism (e.g. Carbohydrate metabolism, Amino acid metabolism), Environmental Information Processing (e.g. Membrane transport), and Genetic Information Processing (e.g. Translation) (Figure 4A). Samples were clustered based on Bray-Curtis distances and gene abundances (KEGG level-1), and control samples were closely clustered (Figure 4B). The top-35 pathways with highest gene abundance were identified (KEGG level-2, Figure 4C). Compared with CDnD and control samples, CDpD samples showed significant higher abundances in “Cellular community”, “Excretory system”, “Circulatory system”, “Transport and catabolism”, “Cardiovascular diseases”, “Endocrine and metabolic diseases”, “Substance dependence”, and “Nervous system”, as well as significant lower abundances in “Membrane transport”, “Carbohydrate metabolism”, “Lipid metabolism”, “Nucleotide metabolism”, “Replication and repair”, “Translation”, “Cell growth and death”, “Folding, sorting and degradation”, “Metabolism of terpenoids and polyketides”, “Transcription”, “Endocrine system”. The top-35 pathways were utilized to perform clustering analysis and PCA, in which CDpD samples could be clearly separated from controls (Figure 4D). Besides, “Lysine biosynthesis” and “Coenzyme B biosynthesis” were only annotated in CDpD samples when compared with controls.

**Figure 4.**
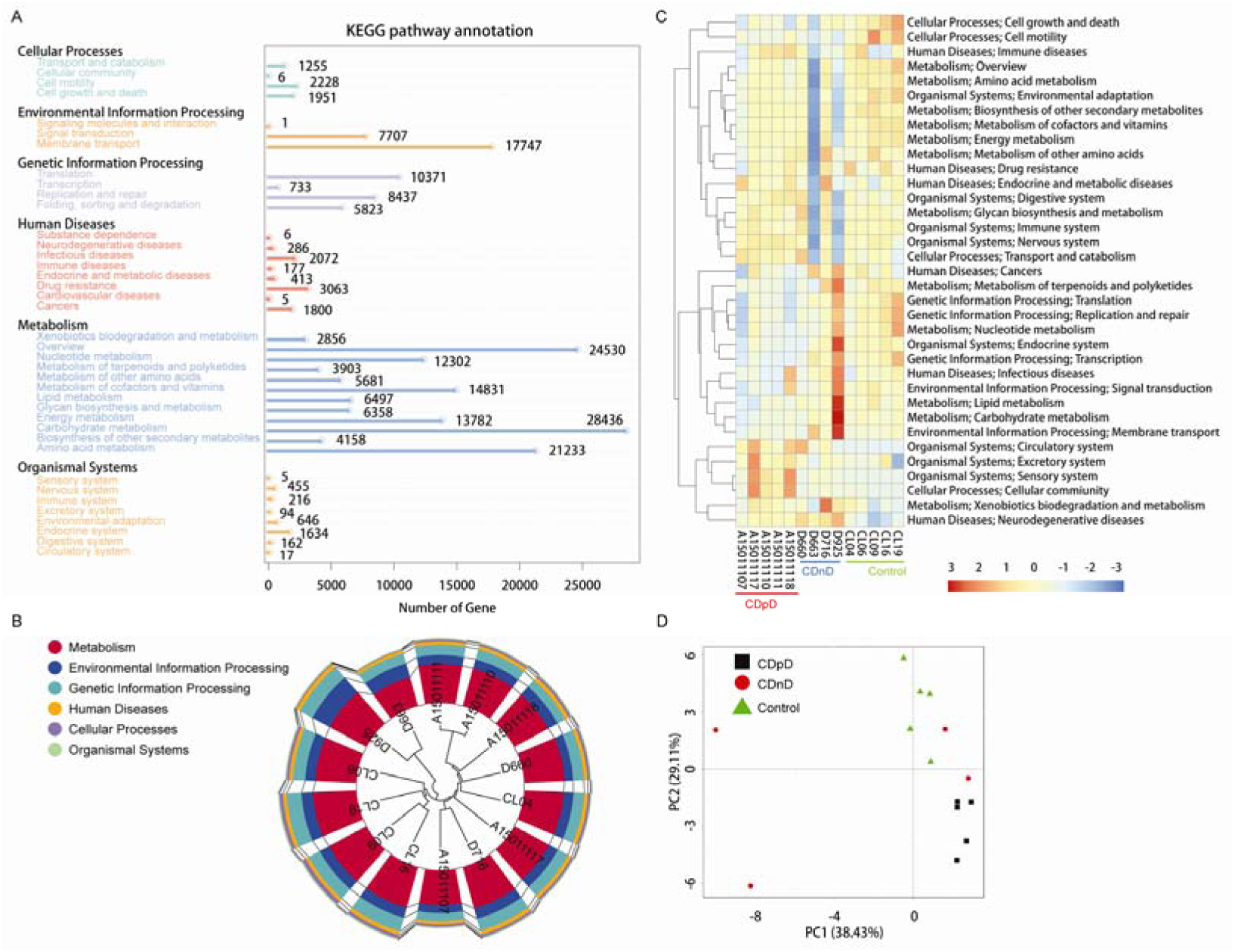
Function annotation of metagenome sequencing data based on KEGG database. (A) Distribution of genes in KEGG pathways. (B) Sample clustering based on Bray-Curtis distances and gene abundances (Level-1). (C) Top-35 pathways (Level-2) with highest gene abundance. (D) PCA based on top-35 pathways (Level-2). PCA: principal component analysis; CDpD: *Clostridium difficile*-positive diarrhea; CDnD: *C. difficile*-negative diarrhea.

### Analysis of A^+^B^+^, A^-^B^+^, and A^-^B^-^ subgroups

#### α-diversity (16S rDNA sequencing data)

In Result-2, Chao1 index, Shannon index, number of observed species in A^+^B^+^ subgroup were much higher than that in A^-^B^+^ and A^-^B^-^ subgroups (Figure 5A-C).

**Figure 5.**
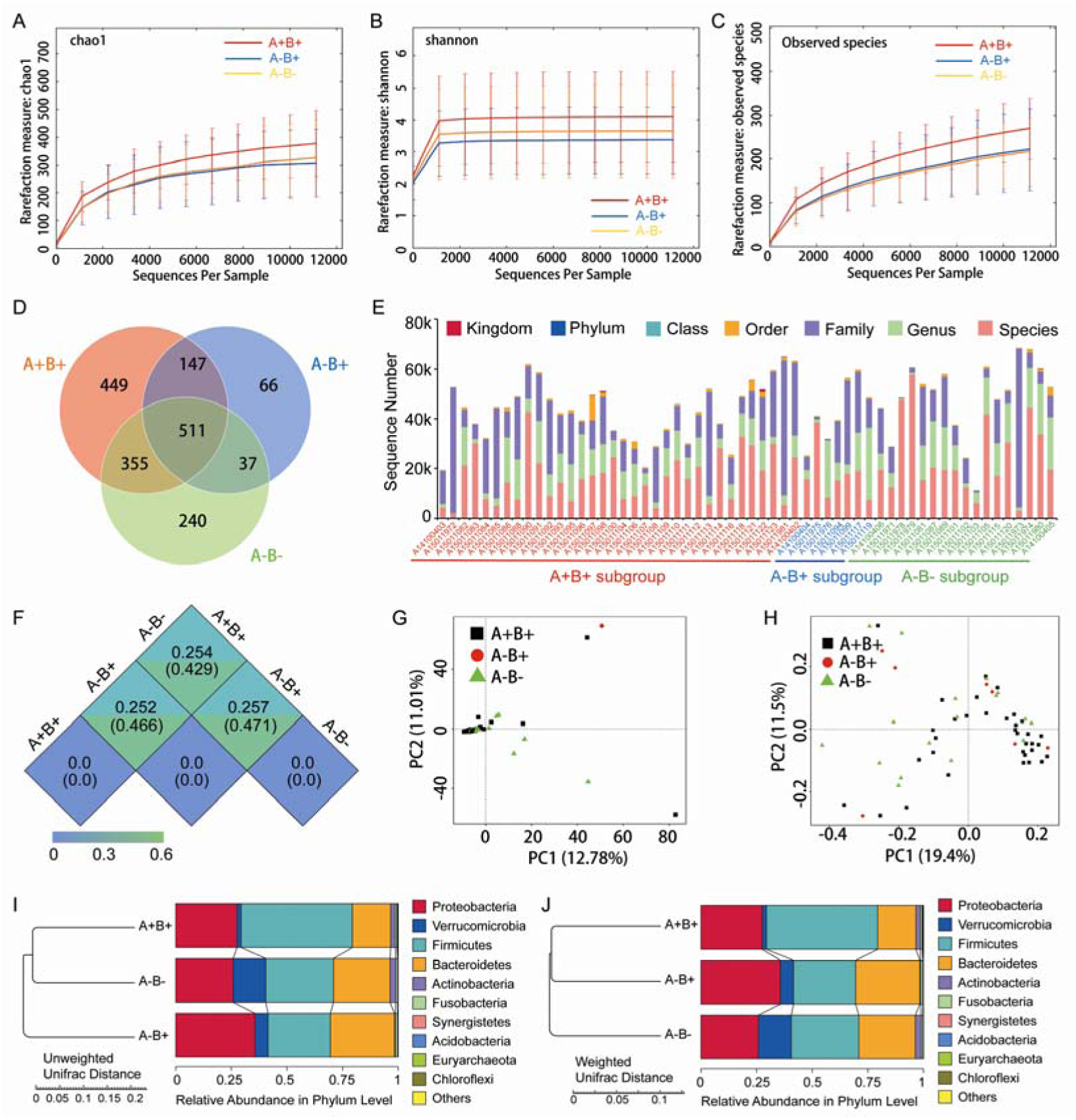
α-diversity and β-diversity of A^+^B^+^, A^-^B^+^, and A^-^B^-^ subgroups based on 16S rDNA sequencing data. (A) Rarefaction-curve based on Chao1 index. (B) Rarefaction-curve based on Shannon index. (C) Rarefaction-curve based on observed species. (D) Venn analysis of OUTs. (E) Distribution of sequences in Kingdom, Phylum, Class, Order, Family, Genus, and Species. (F) Unifrac distance between groups. Upper number: weighted unifrac distance; Lower number: un-weighted unifrac distance. Both numbers represent the index for differences of taxon-diversity between groups. (G) PCA based on OTUs. (H) PCoA based on un-weighted unifrac distance. (I) Sample clustering based on un-weighted unifrac distance and relative abundance at Phylum level. (J) Sample clustering based on weighted unifrac distance and relative abundance at Phylum level. PCA: principal component analysis; OTUs: Operational Taxonomic Units; PCoA: principal co-ordinates analysis.

#### OTUs in each subgroups

In Result-2, 1805 16S rDNA-based OTUs were clustered, among which 511 OTUs were shared by the three groups, whereas 449, 66, and 240 OTUs uniquely presented in A^+^B^+^, A^-^B^+^, and A^-^B^-^ subgroups, respectively (Figure 5D).

#### Taxonomic annotation and β-diversity (16S rDNA sequencing data)

The distribution of sequences in Kingdom, Phylum, Class, Order, Family, Genus, and Species were determined (Figure 5E). The un-weighted unifrac distance (or weighted unifrac distance) were 0.471 (or 0.257), 0.429 (or 0.254), and 0.466 (or 0.252) between A^-^B^+^ and A^-^B^-^, between A^+^B^+^ and A^-^B^-^, and between A^+^B^+^ and A^-^B^+^ (Figure 5F). These results indicated that the differences of taxon-diversity between these subgroups were similar.

In PCA and PCoA based on un-weighted unifrac distance, high inter-sample variability was found across groups, and A^+^B^+^ samples could not be distinguished from A^-^B^+^ or A^-^B^-^ samples (Figure 5G and 5H). However, A^+^B^+^, A^-^B^+^, and A^-^B^-^ subgroups could be distinguished from each other based on un-weighted/weighted unifrac distance and relative abundance at Phylum level (Figure 5I and 5J).

#### Differences of community composition between subgroups

In Result-2, compared with A^-^B^+^ and A^-^B^-^ subgroups, relative abundances of Bacteroidetes, Verrucomicrobia, Bacteroidia, Verrucomicrobiae, Bacteroidales, Verrucomicrobiales, Verrucomicrobiaceae, and *Akkermansia* were lower, whereas relative abundances of Firmicutes, Clostridia, Bacilli, Clostridiales, Lactobacillales, Enterococcaceae, and Ruminococcaceae were higher in A^+^B^+^ subgroup (Figure 6A-D). Moreover, Methylococcaceae, Geobacteraceae, Peptococcaceae, Crenotrichaceae, Mogibacteriaceae, *Acetobacter*, *Dialister*, and *Crenothrix* significantly increased in A^+^B^+^ subgroup in comparison with A^-^B^-^ subgroup. Exiguobacteraceae, Rickettsiaceae, Promicromonosporaceae, Procabacteriaceae, *Providencia*, *Cellulosimicrobium*, *Wolbachia*, and *Saccharopolyspora* significantly decreased in A^+^B^+^ subgroup in comparison with A^-^B^-^ subgroup.

**Figure 6.**
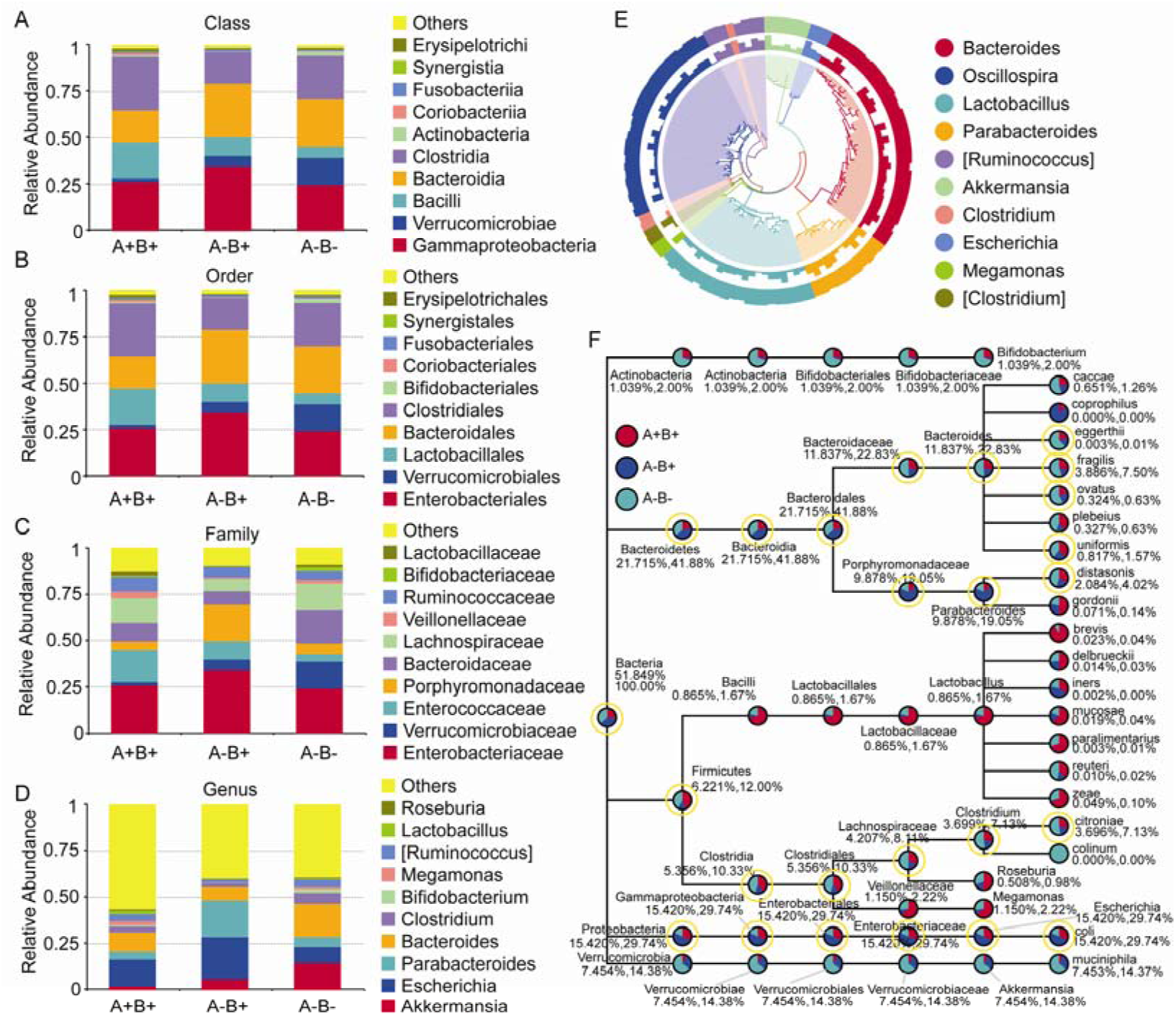
Taxonomic differences and phylogenetic relationships based on 16S rDNA sequencing data in A^+^B^+^, A^-^B^+^, and A^-^B^-^ subgroups. (A) Taxonomic differences between A^+^B^+^, A^-^B^+^, and A^-^B^-^ subgroups (Class-level). (B) Taxonomic differences between A^+^B^+^, A^-^B^+^, and A^-^B^-^ subgroups (Order-level). (C) Taxonomic differences between A^+^B^+^, A^-^B^+^, and A^-^B^-^ subgroups (Family-level). (D) Taxonomic differences between A^+^B^+^, A^-^B^+^, and A^-^B^-^ subgroups (Genus-level). (E) Phylogenetic relationships between OTUs annotated into top 10 Genera. Inner cycle: phylogenetic tree based on representative sequences of OTUs; Middle cycle: relative abundances of OTUs; Outer cycle: reliability of OUT annotation. (F) Taxonomic differences in selected taxa at the levels of Phylum, Class, Order, Family, and Genus. The former number = (mean relative abundance of this taxon in all sample)/(mean relative abundance of all taxa in all sample)×100%; The later number = (mean relative abundance of this taxon in all sample)/(mean relative abundance of selected taxa in all sample)× 100%. Taxon with yellow cycle: this taxon was balanced across sample. OTUs: Operational Taxonomic Units.

#### Phylogenetic relationship

Phylogenetic relationships between the representative sequences of all OTUs corresponding to top 10 taxa in Genus were studied (Figure 6E), and a taxonomic tree was constructed (Figure 6F). *Bacteroides eggerthii*, *Bacteroides fragilis*, *Bacteroides ovatus*, and *Clostridium citroniae* in A^-^B^-^ subgroup were much higher than that in A^-^B^+^ and A^+^B^+^ subgroups. These taxa might be associated with diarrhea. The abundances of Proteobacteria, Gammaproteobacteria, Enterobacteriales, Enterobacteriaceae, *Escherichia*, and *Escherichia coli* in A^-^B^+^ subgroup were much lower than that in A^+^B^+^ and A^-^B^-^ subgroups. These taxa might be associated with A^-^B^+^.

## Discussion

To better understand how microbiome is associated with diarrheal, CDpD, and *C. difficile* type, we characterized the gut microbiomes of healthy individuals and ICU individuals with CDpD or CDnD. Consequently, we found gut microbiota compositions with potential relations with CDpD and *C. difficile* type in ICU patients.

Analyses on Shannon index and observed species indicated that the microbial diversity of CDpD group was lower than that of CDnD and control groups, and this agreed well with previous study in elderly (age□≥□65) hospitalized patients (Milani et al. 2016). The un-weighted unifrac distance between CDpD and control/CDnD was higher than that between CDnD and control. In PCA and PCoA, CDpD samples could be distinguished from CDnD samples and controls, whereas CDnD samples could not be separated from controls. These results indicated that the differences of taxon-diversity between CDpD and control/CDnD were much higher than that between CDnD and control, and that the appearance of *C. difficile* was strongly associated with gut microbiota composition.

Differences of community composition between groups were investigated. As a consequence of metagenome analysis, the abundances of 10 taxa (e.g. Deferribacteres, Cryptomycota, Acetothermia) in CDpD group were significantly higher than that in CDnD group, and based on the 10 significantly different taxa, CDpD samples could be distinguished from CDnD samples in PCA. Reportedly, Deferribacteres is a harmful phylum in intestinal microbiota (Cheng 2016). Generally, Cryptomycota is absent in the healthy human gastrointestinal tract Hallen-Adams H E 2015). Acetothermia can be detected in the gut microbiota of rat rather than human (Khan et al. 2016). Additionally, the abundances of 2 taxa (i.e. Saccharomycetes and Clostridia) were significantly lower in CDpD in comparison with control. *Saccharomyces boulardii* is a non-pathogenic yeast that promotes intestinal immunoglobulin-A immune response to *C. difficile* toxin A and thus protects against recurrent *C. difficile* colitis and diarrhea (Qamar et al. 2001). A balanced gut microbiota is characterized by conserved features like predominance of Bacteroidia and Clostridia, and dysbiosis during episodes of disease generally associates with the rise of un-Bacteroidia or un-Clostridia bacteria (Winter and Baumler 2014). Therefore, CDpD might be associated with the decrease of beneficial taxa (i.e. Saccharomycetes and Clostridia) and the increase of harmful taxa (e.g. Deferribacteres, Cryptomycota, Acetothermia) in gut microbiota. Also, Saccharomycetes and Clostridia might serve as future candidate targets in ICU CDpD patients to help them re-establish healthy gut microbiota. Actually, *Saccharomyces boulardii* has been utilized to prevent antibiotic-associated diarrhea in adult hospitalized patients in clinical trials (Pozzoni et al. 2012; Ehrhardt et al. 2016).

Moreover, based on 16S rDNA sequencing data, relative abundances of some taxa (e.g. Gammaproteobacteria, Enterobacteriaceae, Porphyromonadaceae, and *Akkermansia*) were higher in CDpD group in comparison with CDnD and control groups. Our results were partially consistent with previous study, in which Gammaproteobacteria and Enterobacteriaceae were significantly over-representated in CDI samples, whereas Porphyromonadaceae and *Akkermansia* were significantly under-representated in CDI samples, in comparison with non-CDI samples. Reportedly, *A. muciniphila* is an intestinal representative of the Verrucomicrobia, and it associates with health in humans. Its reduced levels are observed in patients with inflammatory bowel diseases, and it possesses health-promoting activities and anti-inflammatory properties (Derrien et al. 2016). Although *Akkermansia* may improve barrier function, its over-representation may be caused by modifications in the gut micro-environment and reflects enteric mucosa inflammation (Zwielehner et al. 2011). Thus, CDpD might also be related with the increase of Gammaproteobacteria and Enterobacteriaceae in gut microbiota.

Furthermore, differences in microbiota pathways between the three groups were studied using metagenome sequencing data. Compared with CDnD and control samples, CDpD samples showed significant higher abundances in pathways like “Excretory system”, “Circulatory system”, “Transport and catabolism”. “Lysine biosynthesis” and “Coenzyme B biosynthesis” were only annotated in CDpD samples in comparison with controls. Therefore, CDpD might be related with these pathways in gut microbiota, and drugs inhibiting these pathways or the colonization of Deferribacteres, Cryptomycota, Acetothermia, Gammaproteobacteria, and Enterobacteriaceae might be effective in limiting *C. difficile* blooming.

In CDpD group, the community richness and microbial diversity of A^+^B^+^ subgroup were higher than that of A^-^B^+^ and A^-^B^-^ subgroups. The similar un-weighted unifrac distance (or weighted unifrac distance) between A^-^B^+^ and A^-^B^-^, between A^+^B^+^ and A^-^B^-^, and between A^+^B^+^ and A^-^B^+^ indicated the similar differences of taxon-diversity between these subgroups. High inter-sample variability was found in PCA and PCoA, and A^+^B^+^ samples could not be distinguished from A^-^B^+^ or A^-^B^-^ samples. Besides, some taxa were significantly different between A^+^B^+^ and A^-^B^-^ subgroups (e.g. Methylococcaceae, Geobacteraceae, Peptococcaceae, Crenotrichaceae, Mogibacteriaceae, *Acetobacter*, *Dialister*, *Crenothrix*, Exiguobacteraceae, Rickettsiaceae, Promicromonosporaceae, Procabacteriaceae, *Providencia*, *Cellulosimicrobium*, *Wolbachia*, *Saccharopolyspora*). These results suggested that *C. difficile* types might slightly relate to gut microbiota composition in ICU patients with CDpD.

However, some limitations were found in this study. In fact, age and antibiotic usage were not comparable among the three groups, since the clinical complexity of ICU patients and the application of polypharmacy did not allow us to identify a sufficient number of samples for each group or to standardize antibiotic treatment.

In ICU patients, the development of CDpD might be associated with alterations in gut microbiota, including decreases in beneficial taxa (i.e. Saccharomycetes and Clostridia) and the increases in Deferribacteres, Cryptomycota, Acetothermia, Gammaproteobacteria, and Enterobacteriaceae. Besides, *C. difficile* types might slightly relate to gut microbiota composition in ICU patients with CDpD. These results might provide a better understanding of the mechanism of CDpD in ICU patients and directions for developing novel therapies to re-construct healthy gut microbiota.

## Methods

### Study population and sample collection

This study was approved by the Ethics Committee of Xiangya Hospital affiliated to Central South University, China (ID 201212027). All participants granted informed consent before enrollment, and all investigations followed the principles of the Declaration of Helsinki.

A total of 112 subjects were enrolled into this study between March 2014 and December 2014. Subjects included non-pregnant healthy individuals and non-pregnant ICU patients with signs of hospital-onset diarrhea, namely, diarrhea occurred 48 hours after hospital admission with stool 3 or more times within 24 hours. Subjects with inflammatory bowel disease (IBD) and diarrhea occurred less than 48 hours after hospital admission were excluded from the study.

A stool specimen of at least 300 μl was obtained from each subject and then frozen at -20°C. For the samples from ICU patients, *C. difficile* was screened within 24 h using 16s rDNA-PCR (PS13: 5’-GGAGGCAGCAGTGGGGAATA-3’, PS14: 5’-TGACGGGCGGTGTGTACAAG-3’), sequencing, and BLAST alignment. Samples with positive results were classified into CDpD group (N=58), and samples with negative results were classified into CDnD group (N=33). Besides, non-diarrheal stool samples were obtained from healthy individuals who did not administer antibiotics within 1 month and without any signs of diarrhea within 7 days before sample collection (N=21). For stool specimens in CDpD, CDnD, and healthy control groups, 16S rDNA V4 sequencing-based microbiota analysis was performed (Result-1). Moreover, 14 samples were selected from these groups (i.e. 5 samples in CDpD group, 4 samples in CDnD group, and 5 samples in control group) to conduct genome sequencing-based metagenomics analysis (Result-3).

According to the PCR results of *C. difficile TcdA* (gene for toxin A; Forward primer: 5’-AGATTCCTATATTTACATGACAATAT-3’, Reverse primer: 5’-GTATCAGGCATAAAGTAATATA CTTT-3’) and *TcdB* (gene for toxin B; NK104: 5’-GTGTAGCAATGAAAGTCCAAGTTTACGC-3’, NK105: 5’-CACTTAGCTCTTTGATTGCTGCACCT-3’), samples in CDpD group were classified into A^+^B^+^ subgroup (N=34), A^-^B^+^ subgroup (N=7), and A^-^B^-^ subgroup (N=17). For stool specimens in A^+^B^+^, A^-^B^+^, and A^-^B^-^ subgroups, 16S rDNA V4 sequencing-based microbiota analysis was performed (Result-2).

### Clinical data collection

For each subject, data about age, gender, duration of hospital stay, *C. difficile* detection, other diseases (i.e. diabetes, cancer, hematopathy, respiratory failure, renal insufficiency, and tuberculosis), surgery, and exposure to immunosuppressant, glucocorticoids, and antibiotics were collected.

### Amplification and sequencing of 16S rDNA

As previously described (Schubert et al. 2014; Milani et al. 2016), DNA was extracted from stool samples, and the V4-region (primers: 515F and 806R) of 16S rDNA were amplified and sequenced using Illumina HiSeq2500 PE250 Paired-end sequencer.

### 16S rDNA-based microbiota analysis

#### Operational Taxonomic Unit (OTU) clustering

After data preprocessing, UPARSE (version: 7.0.1001, http://drive5.com/uparse/) (Edgar 2013) was used to cluster effective tags into different OTUs (identity ≥ 97%). The tag with highest frequency in a certain OTU was considered as the representative sequence of this OTU.

#### Taxonomic assignment

Taxonomic assignments were performed using the Ribosomal Database Project (RDP) classifier (version: 2.2, http://sourceforge.net/projects/rdp-classifier/) (Wang et al. 2007) and GreenGene database (http://greengenes.lbl.gov/cgi-bin/nph-index.cgi) (DeSantis et al. 2006) with a confidence threshold of 0.8-1. Community composition was analyzed at the levels of Kingdom, Phylum, Class, Order, Family, Genus, and Species, respectively.

#### Phylogenetic relationship

MUSCLE (version: 3.8.31, http://www.drive5.com/muscle/) (Edgar 2004) was utilized to investigate the phylogenetic relationships between the representative sequences of OTUs via conducting multi-sequence alignment.

#### Analysis of α-diversity

Normalization was performed between all samples based on the sample with least data. Then, QIIME (version: 1.7.0) was utilized to analyze α-diversity. Community richness and diversity within community were assessed using rarefaction-curve, Chao estimator (http://scikit-bio.org/docs/latest/generated/generated/skbio.diversity.alpha.chao1.html#skbio.diversity.alpha.chao1, a community richness indicator), Shannon index (http://scikit-bio.org/docs/latest/generated/generated/skbio.diversity.alpha.shannon.html#skbio.diversity.alpha.shannon, a community diversity indicator). Results were visualized using R software (version: 2.15.3).

#### Analysis of β-diversity

QIIME was utilized to analyze β-diversity (i.e. between-habitat diversity). Differences of community composition between samples were assessed using un-weighted unifrac distance, weighted unifrac distance, sample clustering tree, principal component analysis (PCA; tool: FactoMineR package and ggplot2 package in R software), principal co-ordinates analysis (PCoA; tool: WGCNA, stats, and ggplot2 packages in R software).

### Genome sequencing

Total metagenomic DNA was isolated from each stool sample (Milani et al. 2016). After repairing sequence end, adding poly-A, adding sequencing adapters, purification, and PCR, DNA libraries were obtained and then sequenced using Illumina HiSeq2500 Paired-end sequencer.

#### Analysis of metagenomic data

##### Metagenome assembly

After data preprocessing, clean reads with high-quality were *de novo* assembled into scaffolds using SOAP denovo software (version: 2.21, http://soap.genomics.org.cn/soapdenovo.html) (Luo et al. 2012) with parameters of -d 1, -M 3, -R, -u, -F (k-mer length was set as 49, 55, or 59, and only the k-mer with maximum N50 was finally used). Scaffolds were broken into scaftigs via deleting N bases. Then, SoapAligner software was used to align clean reads with these scaftigs (alignment parameters: -u, -2, -m 200), and the reads which did not match these scaftigs were extracted from every sample and pooled together to identify information of rare species. Pooled-reads were also assembled into scaffolds (k-mer length = 55), which were broken into scaftigs. Scaftigs from each sample and pooled-reads were filtered, and scaftigs with a length of ≥ 500 bp were used for statistics and gene calling.

##### Gene calling

Open reading frames (ORFs) were predicted using scaftigs (≥500 bp) and MetaGeneMark software (version: 2.10, http://exon.gatech.edu/GeneMark/meta_gmhmmp.cgi) (W. 2010), and ORFs with length <100 nt were removed. Then, a non-redundant gene catalogue was obtained using CD-HIT software (version: 4.5.8, http://www.bioinformatics.org/cd-hit/, parameters: sequence identity ≥0.95, coverage ≥0.9, -c 0.95, -G 0, -aS 0.9, -g 1, -d 0) (Li and Godzik 2006). Thereafter, SoapAligner software was utilized to align clean reads with this gene catalogue (alignment parameters: -m 200, -x 400, identity ≥ 95%), and genes matched with >2 reads were defined as unigenes. Abundance of each gene in each sample was assessed based on the number of matched reads (r) and gene length (L):

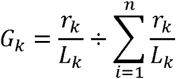

Gene abundance was used to perform core-pan gene analysis, sample correlation analysis, and Venn analysis of gene count.

##### Alignment of unigenes to reference genomes and taxonomical analysis

DIAMOND software (Buchfink et al. 2015) was used to align the metagenomic unigenes to microbial reference genomes of bacteria, fungi, archaea, and viruses in the NR database (version: 2014-10-19) of the National Center for Biological Information (NCBI) (parameters: e-value for BLASTp ≤ 1×10^-5^). For each unigene, only the mapping results with e-value ≤ 10 × minimum e-value were remained, and LCA algorithm in MEGAN software (Huson et al. 2011) was utilized to identify the minimum taxonomic rank without dissent caused by different mapping results.

The abundance of a certain taxon (kingdom, phylum, class, order, family, genus, or species) in a sample was defined as a sum of the abundances of all genes on its member genomes in the sample. The abundances of taxa in every sample were utilized to conduct Krnoa analysis, top-taxa display, sample-taxon clustering, PCA, sample clustering based on Bray-Curtis distances among gene abundances, taxon difference analysis (tool: Metastats (Paulson et al. 2011)), etc.

##### Pathway annotation

To annotate genes, DIAMOND (Buchfink et al. 2015) was used to align unigenes with Kyoto Encyclopedia of Genes and Genomes (KEGG) database (Kanehisa et al. 2014) (parameters: e-value for BLASTp ≤ 1×10^-5^), and results with one HSP > 60 bits were remained.

The counts of annotated genes and the abundances of functions in every sample were utilized to conduct sample-pathway clustering, top-functions display, PCA based on function abundances, sample clustering based on Bray-Curtis distances among function abundances, function difference analysis (tool: Metastats (Paulson et al. 2011)), KEGG pathway display, etc.

### Statistical analyses

Statistical analyses were performed using SPSS 19.0 software to determine the differences of demographic and clinical characteristics among the three groups (i.e. CDpD, CDnD, and control groups), as well as the three subgroups (i.e. A^+^B^+^, A^-^B^+^, and A^-^B^-^ subgroups). For continuous variables, comparison among three groups was performed using one-way ANOVA, whereas comparison between two groups was conducted using independent-samples t-test. In one-way ANOVA, least-significant-difference test and Dunnett’s T3 method were used under the conditions of equal variance and unequal variance, respectively. P-value <0.05 was set as the criterion for statistical difference. For categorical variables, Fisher’s exact test and Pearson’s chi-square test was performed.

## Data Access

In this study, the sequencing data are newly generated data, and they have not been submitted to any public database.

The following software and databases were utilized to analyze the sequencing data: FLASH (http://ccb.jhu.edu/software/FLASH/), QIIME (http://qiime.org/scripts/split_librariesfastq.html), Gold database (http://drive5.com/uchime/uchime_download.html), UCHIME Algorithm (http://www.drive5.com/usearch/manual/uchime_algo.html), UPARSE (http://drive5.com/uparse/), RDP classifier (http://sourceforge.net/projects/rdp-classifier/), GreenGene database (http://greengenes.lbl.gov/cgi-bin/nph-index.cgi), MUSCLE (http://www.drive5.com/muscle/), SOAPaligner soap2.21 software (http://soap.genomics.org.cn/soapaligner.html), SOAP denovo software (http://soap.genomics.org.cn/soapdenovo.html), MetaGeneMark software (http://exon.gatech.edu/GeneMark/meta_gmhmmp.cgi), CD-HIT software (http://www.bioinformatics.org/cd-hit/), etc. Details were shown in **Methods**.

## Acknowledgement

This work was supported by the National Natural Science Foundation of China (No.81601803). Research Fund of Hunan Provincial Health and Family Planning Commission (No. B2016107), Young Scientists Fund of Xiangya Hospital (2014Q05), and Xiangya Sinobio way Health Research Fund (No. xywm2015I11).

## Disclosure Declaration

### Authors’ contributions

AW, JD and CL conceived and designed the experiments. JD, SL and XM performed the experiments. JD, PZ and QD analyzed the data. CL and AW wrote the paper. All authors read and approved the final manuscript.

